# Distinct ensembles in the noradrenergic locus coeruleus evoke diverse cortical states

**DOI:** 10.1101/2020.03.30.015354

**Authors:** Shahryar Noei, Ioannis S. Zouridis, Nikos K. Logothetis, Stefano Panzeri, Nelson K. Totah

## Abstract

The noradrenergic locus coeruleus (LC) is a crucial controller of brain and behavioral states. Activating LC neurons synchronously *en masse* by electrical or optogenetic stimulation promotes a stereotypical “activated” high-frequency cortical state. However, it has been recently reported that spontaneous LC cell-pairs have sparse yet structured time-averaged cross-correlations, which is unlike the high synchrony of *en masse* neuronal stimulation. This suggests the untested possibility that LC population activity may be made of distinct multi-cell ensembles each with unique temporal evolution of activity. We used non-negative matrix factorization (NMF) to analyze large populations of LC single units simultaneously recorded in the rat LC. Synthetic spike train simulations showed that NMF, unlike the traditional time-averaged pairwise correlations, detects both the precise neuronal composition and the activation time courses of each ensemble. NMF identified the existence of robust ensembles of spontaneously co-active LC neurons. Since LC neurons selectively project to specific forebrain regions, we hypothesized that individual LC ensembles produce different cortical states. To test this hypothesis, we triggered local field potentials (LFP) in cortical area 24a on the activation of distinct LC ensembles. We found four cortical states, each with different spectro-temporal LFP characteristics, that were robust across sessions and animals. While some LC ensembles triggered the activated state, others were associated with a beta oscillation-specific state or a reduced high frequency oscillation state. Thus – in contrast to the stereotypical “activated” brain state evoked by *en masse* LC stimulation – spontaneous activation of distinct LC ensembles can control a multitude of cortical states.

## Introduction

Flexible behavior is associated with transitions across diverse cortical states. For example, various states of wakefulness, perceptual ability, and behavioral activity are associated with different cortical local field potential (LFP) and EEG states each with its own clear spectro-temporal pattern of neural oscillations (1–3). Behavioral state transitions, such as waking from sleep or entering a state of heightened stress and reacting more quickly to stimuli, are associated with cortical state transition. These changes are not necessarily driven by external stimuli. Instead, cortical state can be controlled by factors internal to the organism (e.g., sleep need, perceived stress) and therefore arise from self-organized neuronal interactions. It remains unclear exactly which interactions among neurons control cortical states.

Cortical states are mediated, at least in part, by the noradrenergic brainstem nucleus, locus coeruleus (LC). The LC releases norepinephrine to modulate neuronal excitability (4–7). Noradrenergic neuromodulation of cortical state has been studied using direct electrical or optogenetic stimulation. Such stimulation evokes highly synchronous and *en masse* activation of many LC neurons because this brainstem nucleus (in rats) contains ∼1,600 neurons tightly packed into a small volume of ∼200 × 500 × 1000 um (8, 9). For instance, a low current (0.03 - 0.05 mA) pulse evokes spiking up to 400 um from stimulation site in rat LC (10). Thus, even minimal levels of stimulation will synchronously activate many LC neurons given the small dimensions of the nucleus. Such *en masse* LC population activation evokes a stereotypical “activated” cortical state characterized by increased high-frequency oscillation power, regardless of whether the subject is anesthetized or non-anesthetized (10–13). Critically, it is still unknown how spontaneous population activity in the LC, as opposed to *en masse* LC activity evoked by stimulation, relates to the control of cortical state.

Although modulation of cortical state by the LC has largely been studied using external artificial stimulation of LC, cortical state emerges from spontaneously occurring, internal neuronal interactions. The spontaneous activity of the LC neuronal population has been traditionally thought to be highly synchronous (14–19), akin to the *en masse* neuronal population activity evoked by LC stimulation. However, recent findings suggest that this standard view might not describe spontaneous LC activity accurately. Graph-theoretic analysis of time-averaged cross-correlations among pairs of spontaneously active LC neurons in anesthetized rats has reported sparse yet structured pairwise correlations that are unlike the high synchrony of *en masse* stimulation (20). These recent findings suggest that LC population activity might *potentially* consist of multi-cell ensembles that are activated at different times. Importantly, however, these prior pairwise graph-theoretic analyses were based upon time-averaged measures (20) and were therefore unable to detect multi-cell ensembles or to resolve ensemble activity over time. Here, we used non-negative matrix factorization (NMF) to analyze large populations of simultaneously recorded single units in the rat LC. Synthetic spike trains simulations showed that NMF, unlike the traditionally used time-averaged pairwise correlations, detects both the precise neuronal composition and the activation time courses of each ensemble. This allowed us to demonstrate that LC population activity consists of discrete LC ensembles each with its own evolution of activity over time.

Given that LC neurons selectively project to specific forebrain regions, we investigated how individual LC ensembles produce different cortical states. One possibility is that different LC ensembles may simply evoke the stereotypical activated state (the only cortical state associated with LC neuromodulation of the forebrain), as observed in prior anesthetized and non-anesthetized stimulation experiments, but with different LC ensembles being associated with variations in cortical activated state duration or power magnitude. A second and more intriguing possibility is that distinct LC ensembles evoke different cortical states, each with a different spectral signature. This latter case would support a new perspective that LC ensembles may provide a nuanced contribution to the broad set of cortical states that characterize flexible behavior.

Since our methods allowed tracking the spontaneous temporal dynamics of individual LC ensembles, we could relate ongoing LC ensemble dynamics to cortical state dynamics. By using the time-dependent power of the simultaneously recorded LFP in cortical area 24a as an index of cortical state, we performed LC ensemble-triggered LFP (LCET-LFP) analysis, which triggered cortical state on the activation of each distinct LC ensemble (21, 22). In contrast to the standard view that LC population activity evokes a stereotypical activated cortical state, we observed heterogenous cortical states with different spectral and temporal properties that depended on which LC ensemble was active. Importantly, when different ensembles were spontaneously coactivated (more similar to *en masse* synchrony of population activity), the associated cortical states were more homogenous and were more similar to the stereotypical activated state resulting from electrical or optogenetic stimulation. Our results detail spontaneously occurring brain-internal neuronal dynamics that control cortical state by demonstrating that the spontaneous structuring of LC population activity as a collection of discrete ensembles can evoke different cortical states.

## Results

We aimed first to establish whether spontaneous LC population activity was made of subsets of coactive neurons (ensembles) activating spontaneously at different times and then, if so, to characterize their relationship with ongoing cortical state using LCET-LFP. We recorded many LC single units simultaneously (range: 5 to 34 units; average: 19 units; N = 15 male rats) using a silicon probe with 32 electrodes confined to the core of the LC nucleus. Probe location was verified histologically in coronal tissue sections. Neuronal identity was confirmed at the end of the experiments using intra-peritoneal injection of the alpha-2 agonist, clonidine, which inhibited spiking on all electrodes. Spikes recorded from outside the LC core would not have been inhibited due to the lack of alpha-2 adrenergic receptors in nearby brain structures (21). We simultaneously recorded cortical local field potential (8 kHz lowpass filtered) from cortical area 24a (anterior cingulate cortex) (22) using a tungsten electrode in 9 of the 15 rats.

### Spontaneous LC population activity consists of distinct ensembles with independent temporal dynamics

It is currently unknown whether spontaneous LC population activity is made of ensembles (i.e., sub-sets of simultaneously coactive neurons) and how the activity of ensembles changes over time. We assessed whether LC population activity consists of ensembles using non-negative matrix factorization (NMF) on the population vector made of the spike counts of simultaneously recorded single units, independently for each rat. To create the spike count population vectors, we binned activity in sliding windows that were 100 msec long, which is the time scale capturing most of the synchrony among LC single unit pairs (20). **Figure 1A** illustrates how NMF works on *hypothetical* single unit spiking data. NMF decomposes the matrix containing the population vectors at all time points as a sum of K non-negative *spatial modules*, each multiplied by a non-negative *activation coefficient*. A spatial module may be thought of as a specific, often-recurring, population firing pattern. Formally, it is a vector specifying the relative strength of firing of each single unit within the population (23, 24). Thresholding these spatial module values defines the specific single units that were significantly active within each module. On the other hand, the activation coefficients of each spatial module at any given time describe how strongly the spatial module is recruited at that time. We determined K for each rat based on two criteria. First, the chosen K explained a high amount of variance in the data with the fewest possible number of spatial modules, K. In other words, K was in the “elbow” region of the reconstruction error, which when plotted as a function of the possible number of spatial modules, meant that a higher K would have given diminishing returns in terms of data reconstruction accuracy. Second, the selected value for K yielded a stable recovery of the spatial modules from the data regardless of the random initialization of the decomposition optimization procedure (see Methods for additional details and **Figure S1**). By thresholding the time course of activation coefficients to distinguish the times of significant recruitment of each spatial module, we defined the times of activation of each spatial module. The spatial modules will be referred to as “ensembles” and the activation times of spatial modules will be referred to as “ensemble activation times.” In order to study the contribution of LC population activity to cortical state, in the spirit of prior work (25), after using NMF to isolate LC ensembles we then performed a LCET-LFP analysis. In this analysis, we triggered the cortical LFP on localized neuronal events – specifically the activation of distinct LC ensembles (**Figure 1A**).

**Figure 1.**
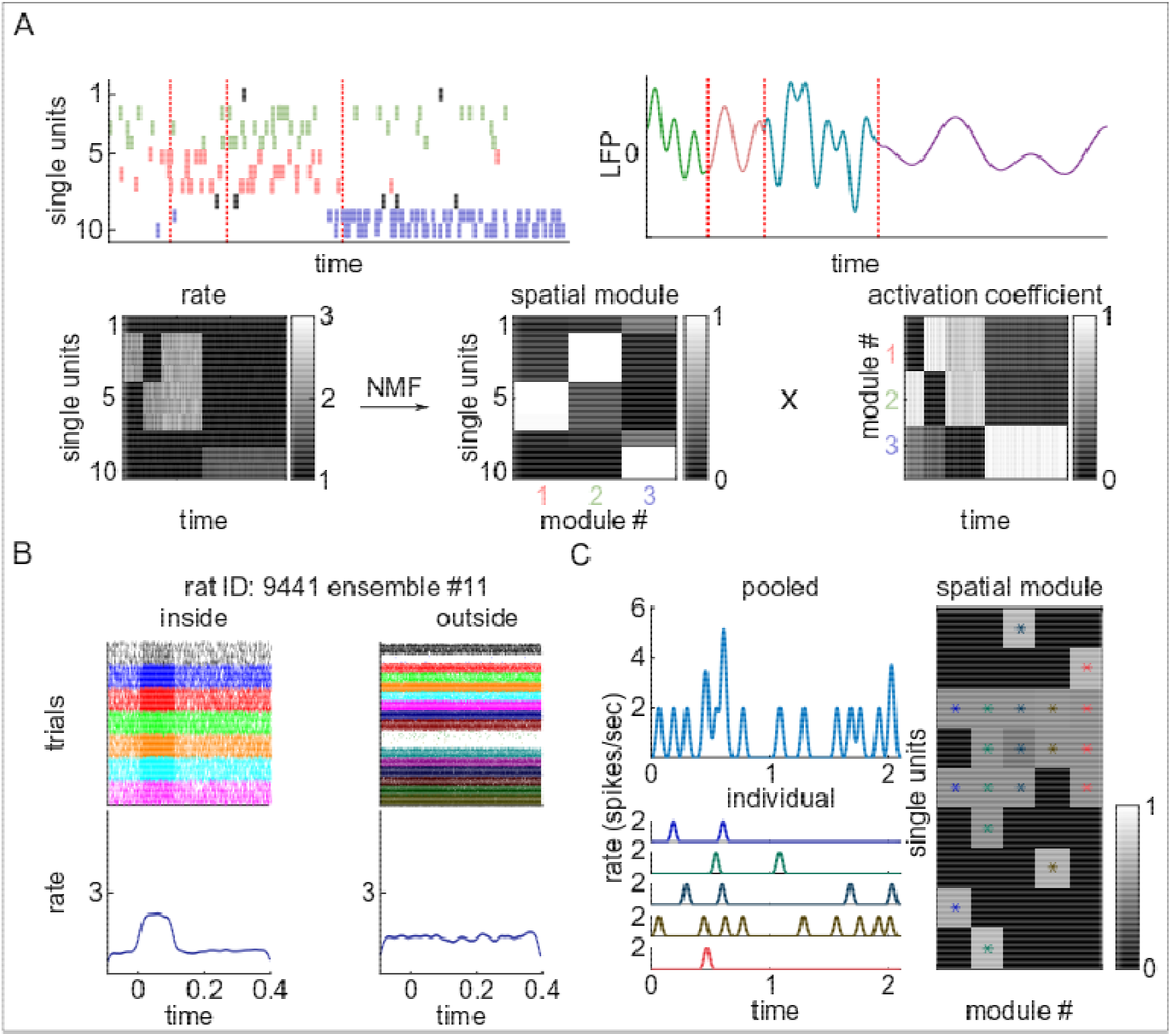
Spontaneous LC neuronal population activity consists of discrete ensembles. **(A)** An example using *hypothetical data* to illustrate how NMF detects ensembles and the time courses of ensemble activity. The spike rasters of 10 single units along with an illustration of cortical LFP is depicted in the top panel. The spikes of different single units, which co-activate as a group and therefore belong to various ensembles, are plotted in distinct colors. The different cortical LFP states are also plotted in various colors. The bottom left panel illustrates the spike rasters of the neuronal population as binned spike rate over time. Larger spike rates are in lighter color. The bottom right two plots show the outcome of NMF on the matrix of population spike rate. The population rate is decomposed as a sum of K spatial modules (in this example, there are 3). It is apparent from the spike rasters (top left) and their replotting as binned spike rates (lower left), that units 2, 3, and 4 often spike together in the first half of the time window (green spikes). Also, during this time window, units 5, 6, and 7 also spike together (red spikes). In the latter half of the window, units 9 and 10 tend to spike together (blue spikes). Therefore, there are 3 ensembles. Note that the first two ensembles are also co-activated briefly just prior to the third ensemble becoming active. The spatial modules (lower middle) resulting from NMF identify 3 specific reoccurring population firing patterns across single units. A threshold is applied to the values of the spatial modules to specify which single units are active in each population firing pattern. For example, module 2 (green) has high values for units 2, 3, and 4; thus, these units are co-active (i.e., ensemble 2). NMF also returns an activation coefficient (lower right) for each module, which represents the strength of recruitment of each module (i.e., each distinct population firing pattern) over time. When activation coefficients are high for a given module (lighter colors), that specific population firing pattern is occurring. Accordingly, NMF recapitulates the patterns observable in the spike raster (top left); ensemble 1 (units 2, 3, and 4, green spikes) and ensemble 2 (units 5, 6, and 7, red spikes) are active during the first half of the recording and are briefly co-active as an ensemble-pair before the latter half of the recording, during which ensemble 3 (units 9 and 10, blue spikes) is active. (**B)** The spike rasters and peri-event time histograms (PETHs) are shown for one exemplar LC ensemble (*actual data*). The left panel shows spike rasters of the single units inside the ensemble aligned to the ensemble activation times (t = 0 sec). In these spike rasters, each ensemble activation event is a “trial.” The PETHs of trial-averaged spike rate across all units in the ensemble are shown below the rasters. The right panel depicts the ensemble activation-triggered spiking of single units that were *not* assigned to that ensemble. The plot shows that units inside the ensemble increased their firing rate at ensemble activation times, whereas units not assigned to the ensemble did not change their firing rate in any systematic way. **(C)** An example from *actual LC data* in which NMF found 5 ensembles among 9 single units. The upper left panel plots the population activity (i.e., the summed spiking of all simultaneously recorded single units) over a 2 second epoch. The lower left panel plots the activation coefficients of the 5 ensembles. Each ensemble is plotted in a different color. The right panel plots the spatial module values. A threshold was applied to these values to specify which single units were significantly active in each spatial module (i.e., ensemble). Threshold crossings are marked with an asterisk with a different color for each ensemble.

NMF revealed for the first time that LC population activity consists of multiple, discrete ensembles of co-active neurons. We detected 146 ensembles from 15 rats. **Figure 1B** shows the spike rasters of 24 simultaneously recorded single units around the time point at which a subset of 7 single units co-activated as an ensemble. The rasters show that those single units which belong to the ensemble increased their firing rate during ensemble activation, whereas other single units not assigned to that ensemble maintained their ongoing pattern of activity without systematic variations around the time of ensemble activation (at t = 0 sec). **Figure 1C** shows another example in which LC population activity was decomposed into 5 distinct ensembles. The ensembles were active in most cases at different times, but occasionally more than one ensemble was simultaneously active (e.g., brown and red lines at t = 0.5 sec). Reconstructing the total population firing rate as function of time through NMF decomposition (i.e., essentially summing up the activation time courses across the 5 ensembles) returned a close approximation of th pooled population spike rate (blue line in upper left panel). This example illustrates that LC neuronal population activity is not merely only sparsely correlated, as shown in prior work (20), but is actually made of a nuanced sequence of distinct ensembles activating at largely non-overlapping times.

Given that this is the first known demonstration of LC ensembles, we further characterized the spatial-temporal and cell type-specific properties of the ensembles. We first assessed the temporal dynamics of ensemble activations. Most ensembles were only transiently active for 100 msec on average. However, the duration of the inactive periods was highly-variable across ensembles (median ± s.d. = 611±295 msec), such that ensembles were quiet for a wide variety of durations before being briefly active for approximately 100 msec (**Figure 2A**). These findings suggest that distinct ensembles spontaneously activate at largely different times relative to one another. The largely non-overlapping activation times of the ensembles will be systematically demonstrated in a later section.

**Figure 2.**
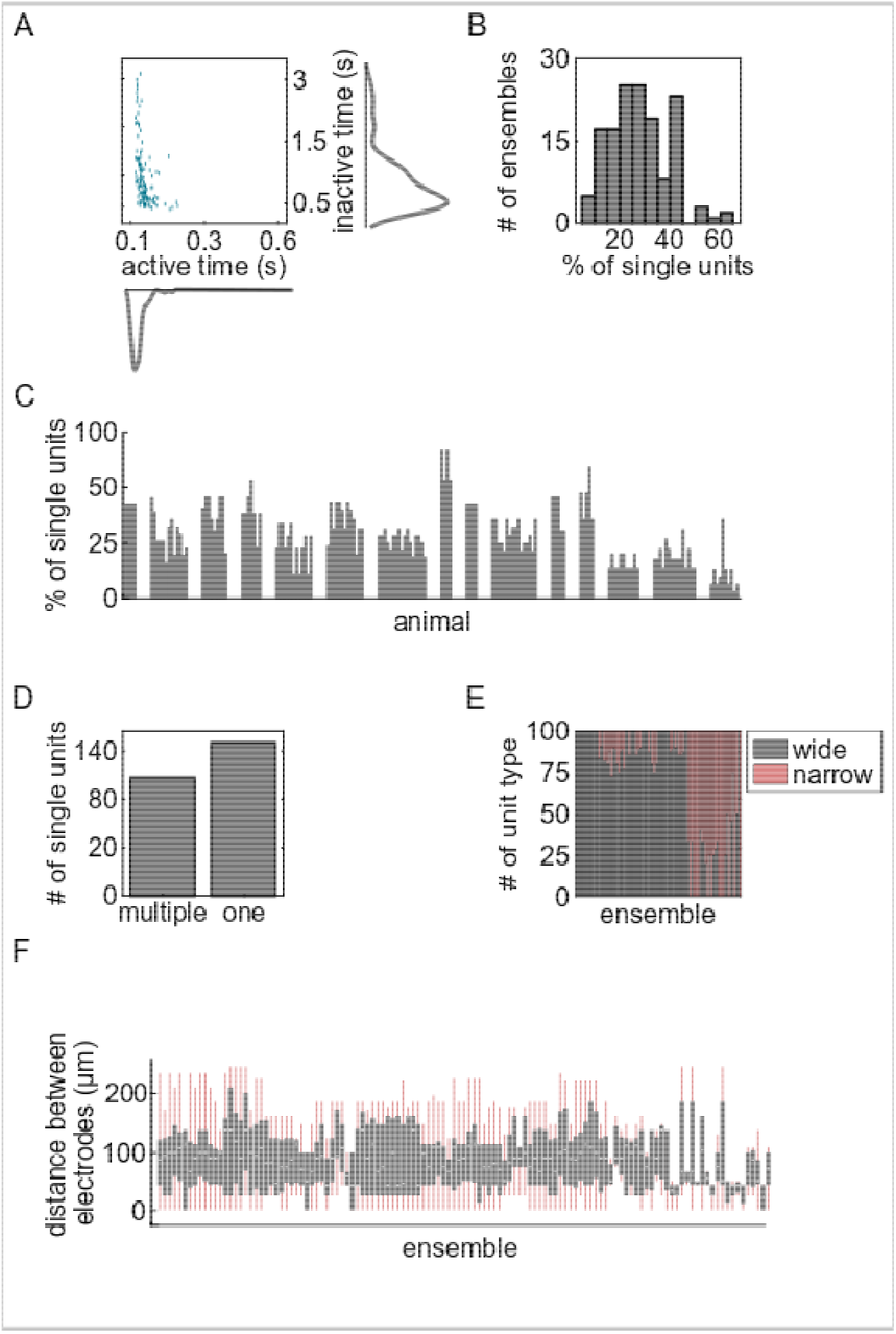
LC ensembles are spatio-temporally sparse and cell type-specific. **(A)** The scatter plot shows the average ensemble active times versus inactive times along with the corresponding histograms. The narrow distribution of active times indicates that most ensembles are only briefly active (∼ 100 msec) before a longer and more variable period of inactivity that is apparent from the broad distribution of inactive times. **(B)** The distribution of ensemble sizes is plotted. The values are the percentage of simultaneously recorded single units that were assigned to an ensemble. On average, each ensemble consisted of 27% of the single units recorded in that experiment. **(C)** Bar plot showing the percentage of single units simultaneously recorded in each rat that belonged to each ensemble. Each bar reports the percentage of one ensemble. Bars are grouped by rat. Note that a single unit can be part of more than one ensemble. **(D)** More single units participated in a single ensemble, but many single units were also observed participating in multiple ensembles. **(E)** The percent of each unit type (wide or narrow spike waveform) making up each ensemble is plotted across ensembles. Rats in which only one single unit type was recorded are not included in this plot. The histogram suggests that ensembles typically consist of one single unit type or at least a majority of one type of unit. **(F)** Boxplots showing the distribution of the distance among the electrodes from which each pair of single units within each ensemble were recorded. Ensembles with only two single units were excluded. Ensembles are spatially diffuse.

We found that ensembles were made of relatively small sub-sets of single units. On average, 27% of single units were active in ensembles relative to the total number of simultaneously recorded single units in each rat (**Figure 2B**). Ensemble size ranged from 6% - 62% of the simultaneously recorded single units (**Figure 2C**). NMF allows single units to participate in more than one ensemble. Therefore, we assessed whether ensembles consisted of totally separate sets of single units or a more complicated structure of overlapping single units. We counted the number of single units assigned to one ensemble, to multiple ensembles, or to no ensemble. Out of 285 single units, 115 single units fired as part of multiple ensembles (40.4%), 149 were active in only a single ensemble (52.3%), and the remaining 21 units did not participate in any ensemble (**Figure 2D**). Although single units did spike in multiple ensembles, the probability that a neuron took part in only one ensemble was higher than the probability that a neuron took part in more than one ensemble (binomial test, p = 0.04).

Our recent work has shown that the LC contains two types of single units, termed “narrow” or “wide” type, which are distinguishable by their extracellular waveform shape and have several physiological and functional differences (20). We assessed (by random resampling) if the proportion of each unit type participating in each ensemble was statistically different from what would be expected if ensembles were formed by units taken randomly regardless of their type. For all rats, the hypothesis that ensembles are formed by combining units regardless of their type was rejected (p < 0.05). Accordingly, ensembles preferentially consist of the same type of unit (**Figure 2E**).

We found that ensembles were made of single units that were spatially diffuse. The location of the maximal average action potential amplitude on the electrode array was used to assign a spatial location to the single unit. We examined the distribution within each ensemble of the distance among the electrodes from which each cell pair within an ensemble was recorded (**Figure 2F)**. There was a wide spread of distances between the units in each ensemble. Therefore, LC ensembles do not follow a topographical arrangement in the dorsal-ventral or medio-lateral aspects of the core of the LC nucleus.

Finally, we assessed how firing strength varied across LC ensembles. In order to quantify the firing strength of LC ensembles, we calculated the average spike rate of all single units within the ensemble (when the ensemble was active) using peri-event time histograms (PETHs). Each event was an ensemble activation time. The PETHs were calculated from 100 msec before each ensemble activation event until 400 msec afterwards. We characterized the diversity of firing strengths across ensembles by clustering the PETHs of 146 ensembles using Principal Component Analysis and Gaussian Mixture Models. When visualizing the data in two dimensions, we observed 3 non-circular masses of data (**Figure S2A**) and, therefore, divided the PETHs into 3 groups. These groups were associated with low, medium, and high changes in spike rate, but had similar activation durations (**Figure S2B**). Most ensembles (88%, green and red in **Figures S2B** and **S2C**) were characterized by a low or medium change in single unit spike rate corresponding to an increase of 1 to 3 spikes per sec (**Figure S2B**). In the maximal case, average spike rate increased by 7 spikes per sec (**Figure S2B**, light purple line), but this was the smallest group of ensembles (**Figure S2C**, light purple). Single unit spike rate for those units within the ensemble was higher when the ensemble was active than when it was inactive (**Figure S2D**, gray, two-sided Wilcoxon rank sum test, Z = 20.9, D = 0.8, power = 0.99, p < 0.001). We also assessed the average spike rate when all single units within an ensemble were merged into a single multi-unit spike train. Again, spike rate within the ensemble was higher during epochs of ensemble activation (**Figure S2E**, gray, two-sided Wilcoxon rank sum test, Z = 14.7, D = 2.6, power = 0.99, p < 0.001). On the other hand, when an ensemble was inactive, multi-unit activity outside of the ensemble was relatively higher (**Figure S2E**, light green, two-sided Wilcoxon rank sum test, Z = 6.8, D = 0.8, power = 0.99, p < 0.001). This is due to those units spiking as members of other ensembles during these epochs. These results show that the firing strength can vary considerably across LC ensembles.

### NMF detects the composition of ensembles and their activation times, whereas graph-theoretic time-averaged pairwise correlations used in prior work does not

Recent work has demonstrated that most pairs of LC single units do not fire in strong synchronization, but rather show sparse yet structured cross-correlations (20). A graph-theoretic community detection analysis on the pairs with significant time-averaged cross-correlations demonstrated cases in which some single units tended to synchronize their spiking with multiple other single units (20). This analysis is suggestive of a sparse structure of interactions among LC neurons, but it is merely compatible with the possibility that spontaneous LC activity is made up of ensembles. Here, we formally demonstrate that graph theory analysis of time-averaged cross-correlations, as used in (20), is neither sufficient for identifying ensembles, nor can it determine the times at which distinct ensembles are active. These limitations are illustrated in the following examples, which are also used to demonstrate the power of NMF to detect distinct ensembles each of which has its own unique temporal evolution of activity.

We generated three different scenarios of simulated spike trains (**Figure 3**). Each scenario consisted of 10 simultaneously recorded single neurons that were all governed by ensemble dynamics. In all three scenarios, the graph-theoretic analysis of cross-correlations shown in the bottom left panels performed exactly as in prior work on LC population activity (20). They revealed the same set of units that were more strongly correlated with each other: units 2 through 7 and units 9 through 10 (bounded by the green and red boxes that denote the two “communities” identified by the graph theoretic analysis). Information about the temporal activation patterns of these communities cannot be obtained from the graph-theoretic analysis because, as in prior work (20), it uses time-averaged neuronal activity. However, the ground-truth ensemble dynamics were radically different in each scenario. In the first and second scenario (**Figure 3A** and **3B**, respectively), spike trains were generated as unique temporal sequences of activation of the two ensembles (ensemble 1 - units 2 through 7; ensemble 2 - units 9 through 10) that were identified by the graph-theoretic correlation analysis as two communities of neurons strongly correlated with each other. However, the graph-theoretic analysis, being based on time-averaged correlations, could not identify that, in these two scenarios, the ensembles had very different activation dynamics. For instance, in the first scenario (**Figure 3A**), ensemble 1 was the only strongly active ensemble for the first part of the simulation and ensemble 2 was the only strongly active ensemble for the latter part of the stimulation. Whereas, in the second scenario (**Figure 3B**), not only were the periods of lone strong activation of each of the ensembles different from the first scenario, but there was also a period at the end of the simulation when there was intermediate co-activation of both ensembles. In both cases, NMF can capture the units that are members of each ensemble (i.e., by thresholding the spatial module values) and it can reconstruct the temporal dynamics of each ensemble both when they are active alone as well as when they are co-active. These two scenarios (**Figures 3A** and **3B**) highlight that the time-averaged graph-theoretic analysis cannot resolve ensemble activation dynamics. Finally, in **Figure 3C**, we considered a third scenario in which the graph-theoretic analysis generated again two communities identical to those in the first two scenarios (Figure **3A** and **3B**). Critically, however, there were actually three distinct ground-truth ensembles: (ensemble 1 - units 2 through 4; ensemble 2 - units 5 through 7; and ensemble 3 - units 9 through 10). Ensembles 1 and 2 had different temporal activation patterns with often only one of the two ensembles being active, but they also had a period in which they were both strongly coactive. The graph theoretic analysis based on time-averaged cross-correlations incorrectly conflated the first two ensembles into a single community made of units 2 through 7. This illustrates that the time-averaged correlation analysis and graph theory detection of communities cannot resolve the identity of individual ensembles. Thus, our prior work (20), which reported the first large-scale single unit recordings in the LC, successfully demonstrated with the time-averaged correlation analysis that LC populations did not fire *en masse*, but this was insufficient to demonstrate whether the LC contains ensembles. Furthermore, it could not measure when each ensemble was active.

**Figure 3.**
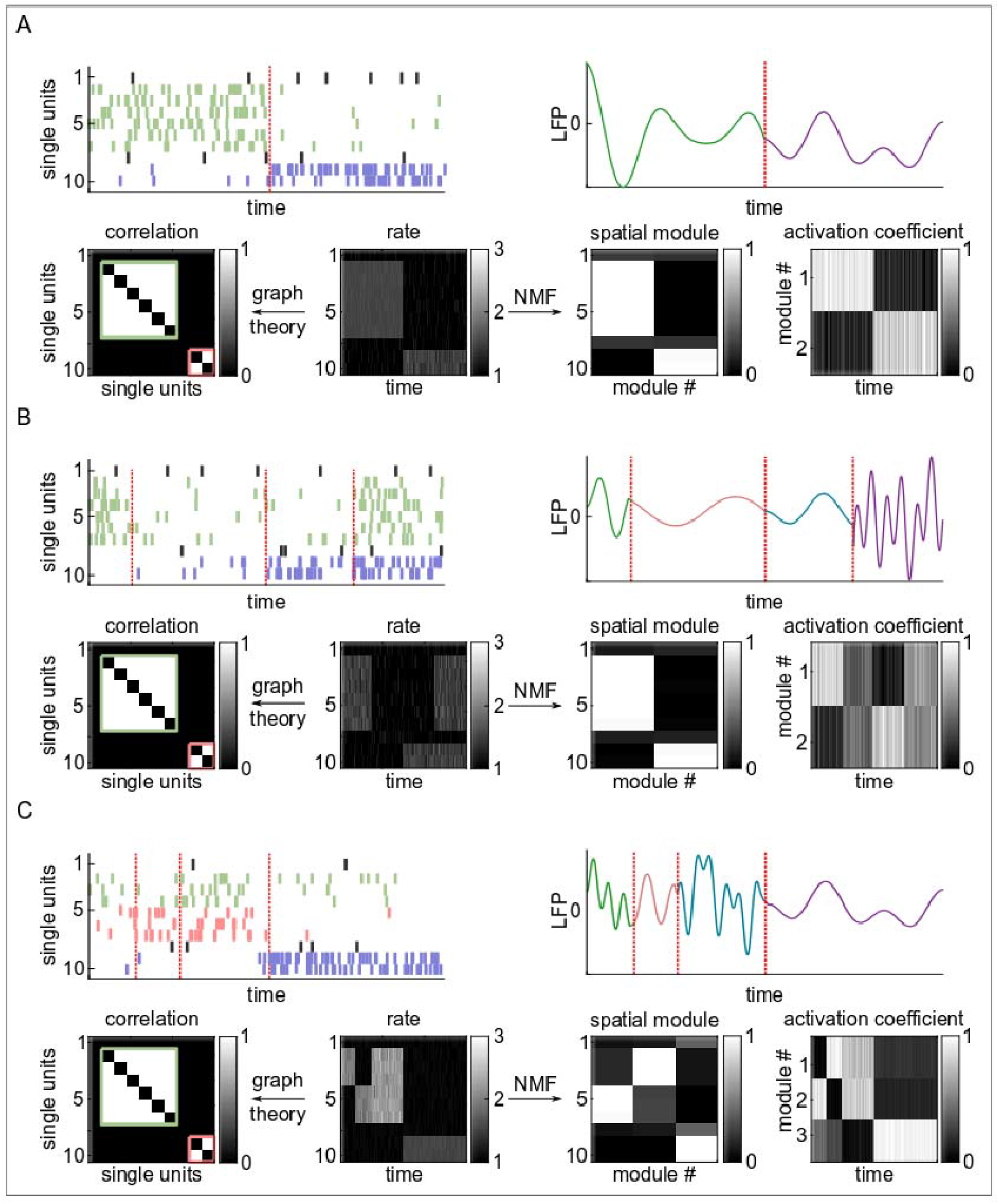
Ensembles cannot be detected using graph-theoretic analysis of time-averaged pairwise correlations. Three different simulated spike rasters were generated in order to compare the NMF method with the graph-theoretic method. In all panels **(A-C)**, the upper left plots depict the generated spike rasters of 10 single units. The spike trains in each panel were generated to have unique ensemble dynamics. The spikes of units belonging to different ensembles are in different colors, while the spikes of units that do not belong to any module are black. These ensembles are ground-truths encoded into the simulation. The red dotted lines indicate transitions in the synthesized LC population activity. The same time points are presented on the top right plots, which show the simulated ongoing cortical activity with different cortical states plotted in different colors. In each panel (**A-C**), the lower panels show the binned spike rates of each unit constructed from the spike rasters (middle), the communities detected by graph theory analysis (left), and the spatial modules (i.e., ensembles) and the ensemble activation time courses (right). In the graph-theoretic analysis, as in prior work (20), the spike count correlation coefficient was calculated from the binned spike counts from the entire recording. Each axis of the lower left plot is a unit and, if the time-averaged correlation for a cell-pair was significant, it was assigned a value of 1 (white) and otherwise 0 (black). Community detection algorithms (as in prior work) were used to detect communities of co-active neurons. These are indicated by the green and red bounding boxes. The result of the NMF analysis (lower right two panels) show the spatial modules and the temporal activation patterns of each ensemble.

Importantly, we verified that the present NMF analysis was able to detect correctly in all 3 scenarios both the identity of the ensembles and the time course of their activation that was used to generate the simulated data (see upper right panels, **Figure 3**). NMF allowed us to go beyond prior work (20) and assess how LC ensemble dynamics related that of other LC ensembles and, most importantly, to ongoing cortical state dynamics.

### LC ensembles are distinct and activate sparsely

Before considering how the activation of LC ensembles relates to cortical states, it is important to further confirm that the NMF-detected LC ensembles are indeed distinct ensembles each with largely non-overlapping activation times, so that different ensembles could potentially produce ensemble-specific cortical states. Our earlier analyses have shown that LC ensembles are activated briefly with long and highly variable duration pauses between one activation and the next (**Figure 1C, 2A**). This result is highly suggestive that the ensembles are distinct because such pauses can maintain largely non-overlapping activation times between ensembles. Here, we more formally characterized these pauses by examining ensemble auto-correlogram and ensemble-pair cross-correlogram troughs. Using the time series of activations of each LC ensemble, we individuated significant troughs in the ensemble auto-correlograms (**Figure 4A**) and the ensemble-pair cross-correlograms (**Figure 4B**). These troughs may be mediated by noradrenergic self-inhibition and lateral inhibition, which is a known property of LC neurons (26–29). A trough in the auto-correlogram, which indicates self-inhibition, occurred in 62% of the ensembles (90 out of 146). For these ensembles, the self-inhibited spiking was most frequent at a 100 msec delay after ensemble activation but could occur as late as 300 msec after ensemble activation (**Figure 4C**). In addition to this self-inhibitory property of LC ensembles, we found that 44% of ensemble-pairs (348 out of 790) had a significant cross-correlogram trough, which indicates a lateral inhibitory interaction. Lateral inhibitory interactions between ensemble-pairs were most frequent after a delay of at ±300 msec, but covered a wide range of variable timings, lasting up to 1 second (**Figure 4D**). These times of self-inhibition and lateral inhibition match well the wide range of pauses apparent in ensemble activity shown in **Figure 2A** and both types of inhibition likely contribute together to generate these pauses in LC ensemble firing. Overall, our results suggest that these inhibitory mechanisms could help produce the sparse activations of distinct LC ensembles, replete with long and highly variable pauses, such that each LC ensemble has its own unique temporal activity dynamics. Critically, the presence of lateral inhibition between ensembles is also key evidence that ensembles are distinct from one another and, in combination with our simulations showing that NMF can discriminate distinct ensembles (**Figure 3**), our analysis of ensemble-pair cross-correlograms strongly supports the claim that NMF can detect distinct ensembles in the LC.

**Figure 4.**
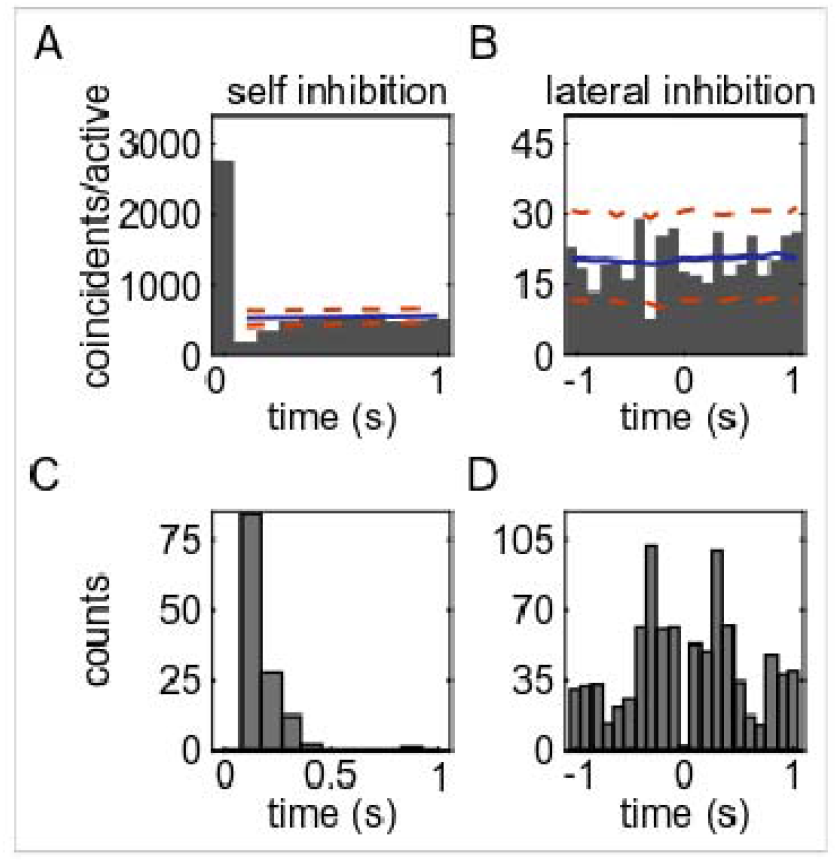
The durations of self-inhibition and lateral inhibition demonstrate that LC ensembles are distinct and their activation pauses occur over 100 milliseconds up to 1 second. **(A)** An example of a self-inhibition of an ensemble visible as a trough in the auto-correlogram. The example is ensemble #5 from rat 2591. The time bin is 100 msec. Significant troughs were defined as those that crossed the lower bound of the 1% pairwise minimum threshold (dashed orange lines), which was calculated from 1000 surrogate data sets constructed by jittering ensemble active times. The solid blue line shows the average of the surrogate correlograms. **(B)** An example of lateral inhibition between an ensemble pair (ensembles 9 and 11 from rat 2632). All details are the same as in panel A. **(C)** The histogram shows the number of significant self-inhibitions across different time bins. For example, the ensemble shown in panel A had significant troughs at 100 msec and 200 msec after it was activated. Therefore, it was counted in the histogram at those two time points. Self-inhibition in almost all cases (98% of all auto-correlogram time points across all ensembles) lasted less than 300 msec. **(D)** The histogram shows the number of significant lateral inhibitions across different time bins. Lateral inhibition peaked at ±300 msec.

### Highly synchronous LC population activity occurs, but rarely

Although our results clearly demonstrate that LC ensembles are distinct from one another and activate with largely non-overlapping time courses that contain numerous highly variable firing pauses, it does not preclude the possibility that LC ensembles co-activate. Examples of this in LC data can be seen in **Figure 1C** (e.g., brown and red lines at t = 0.5 sec). The occurrence of ensemble co-activation is important to quantify because such ensemble population activations would be more like the *en masse* firing evoked in LC stimulation studies that have been used to define the role of the LC in modulating cortical state (10–13).

We quantified the number of ensemble-pairs with significant co-activations. We also assessed the most common delay at which one ensemble was activated after the other in the pair because zero-lag pairwise ensemble activations would be like *en masse* LC activation. Importantly, NMF analysis of synthetic spike trains has demonstrated that this method can detect co-activate ensembles (see **Figure 3B**). We found that 64% of 790 ensemble-pairs had positive cross-correlogram peaks. An example significant positive interaction between two ensembles is shown in **Figure 5A**. We found that most of the significant positive interactions occurred without a delay (**Figure 5B**). Specifically, zero-lag (no delay) interaction was observed in 83% of the 64% of ensemble-pairs with significant positive interactions. In the overall population of 790 ensemble-pairs, this corresponds to synchronous co-activation of 53% of ensembles. Such zero-lag positive interactions indicated some degree of co-activation of ensembles. This result shows that highly synchronous and non-delayed co-activation of LC ensembles occurs among a large proportion of ensemble-pairs. These results suggest that, while LC ensemble are distinct, many ensembles co-activate and can therefore mimic the *en masse* activation of LC neurons due to electrical or optogenetic LC stimulation.

**Figure 5.**
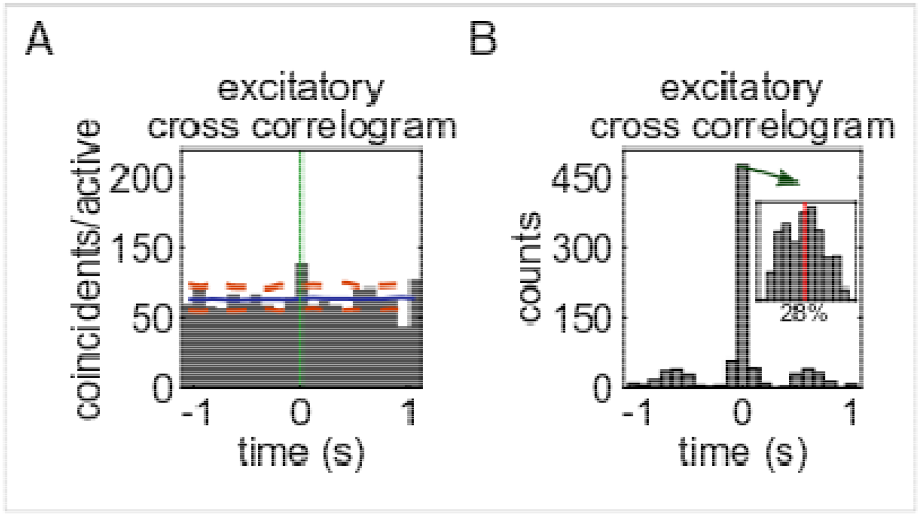
Highly synchronous, zero-lag activation happened among 53% of ensemble-pairs, but its occurrence was rare. **(A)** An example ensemble-pair with significant positive interaction with zero-lag. The activation time of one ensemble in the pair is at t = 0 (green line). The example is between ensembles 1 and 2 from rat 1052. The time bin is 100 msec. Significant peaks were defined as those that crossed the upper bound of the 1% pairwise maximum threshold (dashed orange lines). The solid blue line shows the average of the surrogate correlograms. **(B)** The histogram shows the number of significant ensemble-pair positive interactions across time bins. The majority of positive interactions between ensembles occurs at zero-lag, indicating that many ensemble-pairs synchronously co-activate. The inset presents a histogram of synchrony index values between the ensemble-pairs that had significant zero-lag positive interactions. The average synchrony index is 28%, which indicates that co-activation occurs on only 28% of ensemble activation instances

To measure how often LC ensembles synchronously co-activate, we used a zero-lag synchronization index, which measures what proportion of activation instances of an ensemble were zero-lag co-activations with another ensemble. The calculation was performed on only the 53% of ensemble-pairs with significant zero-lag synchrony. The average synchronization index was 28% (**Figure 5B**, inset), indicating that the 53% of ensemble-pairs with significant zero-lag co-activations were synchronously activated only occasionally (i.e., on 28% of activation instances). Overall, and contrary to the standard view that the LC neuronal population fires *en masse* with a high level of synchrony, these analyses show that LC ensembles had nuanced and largely non-overlapping dynamics with rare ensemble co-activations.

### Different cortical states are associated with activation of distinct LC ensembles

The rich, rarely overlapping dynamics of individual LC ensembles may enable distinct ensembles to evoke different cortical states. Alternatively, all ensembles may promote the stereotypical activated state that is widely viewed as the singular role of noradrenergic neuromodulation of cortical state (1–3, 10–13). We disambiguated between these two possibilities using an LC-ensemble-triggered LFP (LCET-LFP) analysis. The ability to examine the relationship between LC ensemble activity and ongoing cortical state is enabled by NMF, which for the first time has identified LC ensembles and their temporal activation dynamics.

The LCET-LFP analysis triggered cortical area 24a LFP on the activation of individual LC ensembles. NMF identified 89 ensembles in 9 rats from which cortical LFPs and LC population activity were simultaneously recorded. We calculated the LFP spectrogram modulation in a window of 400 msec before ensemble activation until 500 msec afterwards. This window was chosen for two reasons. First, it provided a good tradeoff between temporal and spectral resolution. Second, our previous analyses of cross-correlations and durations of activation and inactivation show that it is unlikely that multiple ensembles were coactive during this window (**Figure 2, Figure 5, and Figure 6**). Therefore, this window ensured that changes in the cortical LFP spectrum were predominantly related to activation of an individual ensemble in the LC. We averaged the spectral modulations for each ensemble over all instances its activation. Visual inspection of the LC ensemble activation-triggered spectra revealed diverse cortical states depending on which ensemble was activated (**Figure S3)**.

**Figure 6.**
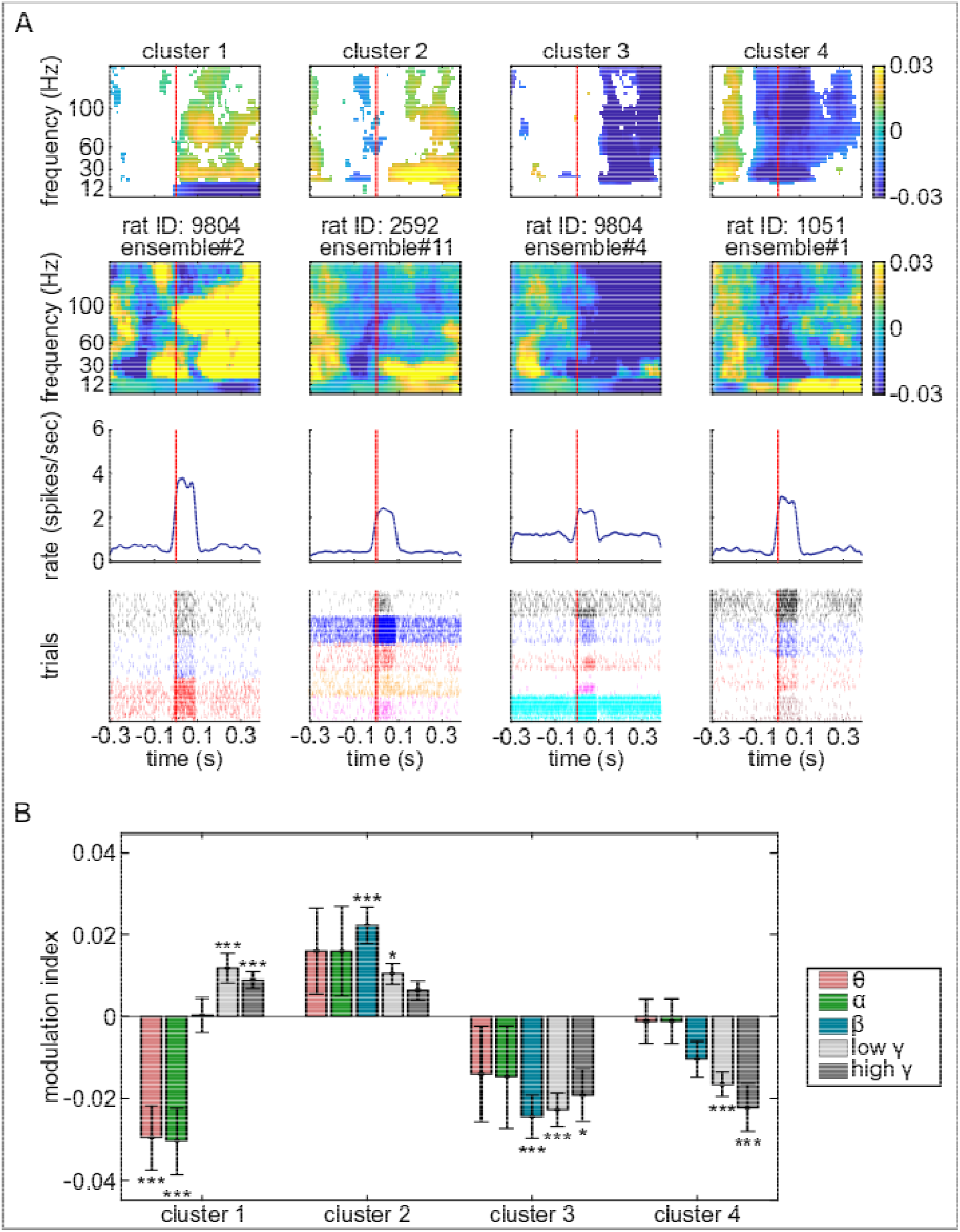
Activation of different LC ensembles are associated with diverse changes in cortical LFP power spectra. **(A)** LFP power spectra were triggered on LC ensemble activation times. The resulting spectra were clustered into 4 types, which are shown as 4 columns. The top row shows the average spectrogram across all spectra in each cluster. Only significant modulations (yellow – increase, blue – decrease) are shown; non-significant values are white. The ensemble activation time is at time 0 and is marked by a solid red line. The lower 3 rows show the activity of an example ensemble from each of the 4 spectral clusters and the ensemble activation-triggered spectrogram for that ensemble. The example spectra show both significant and non-significant values. **(B)** The average modulation indices for different frequency bands are plotted for each cluster. The frequency bands are theta (4-8 Hz), alpha (8-12 Hz), beta (12-30 Hz), low gamma (30-70 Hz) and high gamma (70 - 150 Hz). Significant modulations are assessed by testing the median of the modulation distribution against zero by using Wilcoxon’s signed rank test followed by multiple correction.

Given this apparent diversity of cortical states associated with the activation times of different LC ensembles, we formally assessed if the multitude of LC ensemble-associated cortical states could be consolidated into a few typical patterns. We clustered the spectral modulations associated with each of the 89 LC ensembles. We found 4 predominant types of spectra in the clustering analysis. We chose 4 clusters by first varying the putative number of clusters from 1 to 22 and quantifying the diminishing returns of adding each additional cluster (for details, see Methods and **Figure S4**). Critically, only one of these spectral types (Cluster 1, **Figure 6**) can be described as the stereotypical “activated” cortical state (10–13). The activated cortical state has been previously characterized in extensive prior work that used electrical or optogenetic stimulation to activate LC neurons *en masse* and found that this state is a decrease in theta and alpha band oscillations and an increase in gamma band oscillation power of the mean extracellular field potential in both non-anesthetized and anesthetized preparations (10–13). Our analysis revealed that spontaneous activations of LC ensembles in cluster 1 evoked the activated state (**Figure 6A**). Activation of this sub-set of LC ensembles evoked decreased power in theta and alpha band LFP oscillations and increased power in gamma band oscillations (**Figure 6B**). This stereotypical activated cortical state was associated with activation of 28% of the 89 ensembles.

However, a diverse set of cortical states occurred after activation of different sub-sets of LC ensembles. The second type of ensemble-specific spectra (Cluster 2) was associated with activation of 22.5% of LC ensembles and was characterized by a specific increase in beta oscillations and a small increase in low gamma oscillations, whereas power in other frequency bands did not change significantly. The third type of ensemble-specific spectra (Cluster 3) opposed the direction of the first two spectral types, in that the beta, low gamma, and high gamma bands were decreased. This spectral pattern was associated with 22.5% of the ensembles. In Clusters 1 through 3, the change in cortical state took place after LC ensembles activated. However, the last type of spectrum (Cluster 4) was associated with a change in cortical state that began before LC ensemble activation, namely a decrease in high frequency spectral power. Overall, distinct LC ensembles were associated with spectro-temporally diverse cortical states.

Given that the firing strength was variable across LC ensembles (**Figure S3**), we assessed whether there was a systematic difference in the strength of ensemble population spike rate across the 4 cortical state clusters shown in **Figure 6**. For instance, the stereotypical activated state (cluster 1 cortical spectra) might only occur when an LC ensemble reaches a certain threshold spike rate. Therefore, we assessed the firing strengths of LC ensembles and whether these differences might predict the relationship between which LC ensembles are associated with each of the 4 different cortical states associated with LC ensemble activation. The population spike rate was calculated as the average of all ensemble activation events combined across all single units in the ensemble (i.e., in the spike rasters shown in **Figure 6**, all events of different colors were averaged). The peak of the resulting population spike rate was used to characterize the firing strength of the population in each ensemble. We found that the median population spike rate across ensembles in each cortical spectral cluster differed across clusters (Kruskal-Wallis test, p = 0.0003, *ω*^2^ = 0.9633, *χ*^2^ = 18.82), but post-hoc tests showed that only cluster 1 was different from clusters 2 and 3; therefore, there was no systematic relationship between cortical spectral cluster type and population spike rate (**Figure 7A**). We also examined the peak spike rate of the single units in each ensemble. For this analysis, the spike rates around ensemble activation events were first averaged for each single unit separately (i.e., in the spike rasters shown in **Figure 6**, all events of the same color were first averaged). The peak spike rate of the PETH of each single unit was averaged across units to obtain a measure of single unit firing strength. The median spike rate across all ensembles in each cortical spectral cluster type again differed across clusters (Kruskal-Wallis test, p = 0.0334, = 0.9871, = 8.71). The result was similar to that obtained using the population spike rate, in that the single unit firing rate differed only between clusters 1 and 3 (**Figure 7B**).

**Figure 7.**
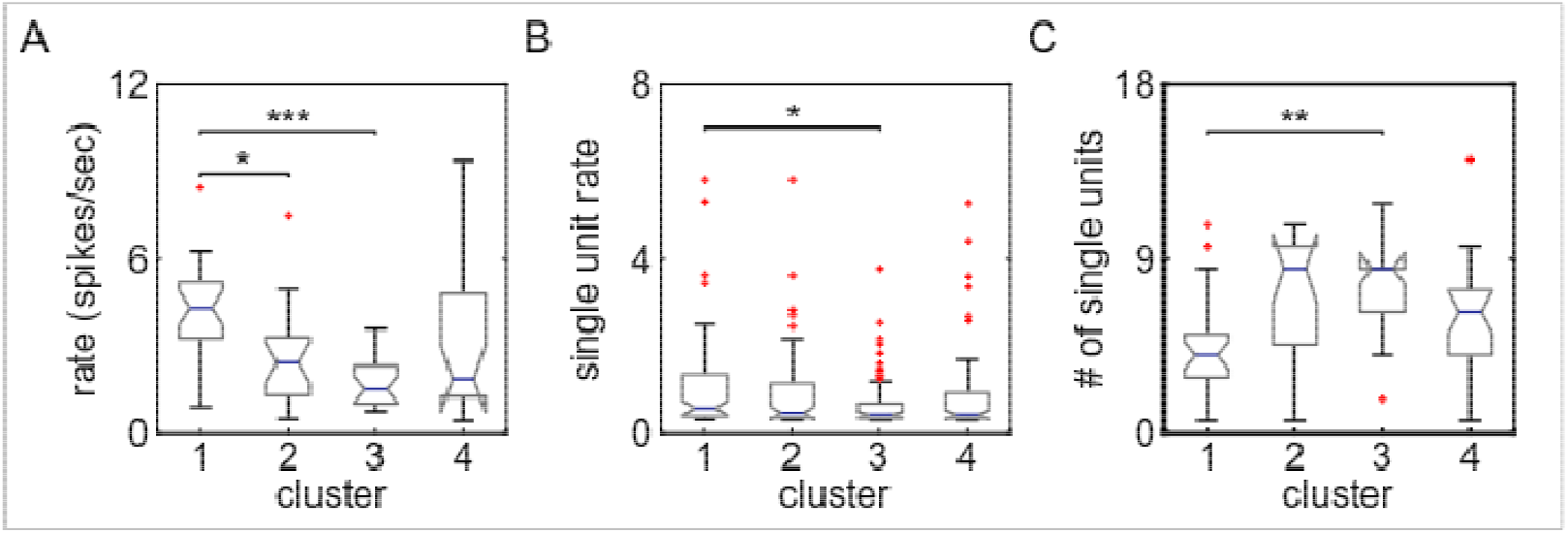
Different cortical states are not related to the intrinsic properties of distinct sub-sets of LC ensembles. **(A)** The box plots show the distributions of maximal spike rates of the ensembles’ PETHs in each spectral cluster. There was a significant difference in spike rate between clusters 1 and 2, as well as between clusters 1 and 3. **(B)** The boxplots illustrate the distribution of the spike rate averaged across single units within the ensembles and separating the ensembles by spectral cluster. A significant difference was observed only between clusters 1 and3. (C) The boxplots show the distributions of the number of single units within the ensembles for the different spectral clusters. A significance difference was found only between spectral clusters 1 and 3.

We also examined another factor that could predict how distinct LC ensembles are associated with different cortical states. Specifically, the size of the ensemble (i.e., the number of single units within the ensemble) might systematically vary with the cortical spectral cluster type. For instance, a type 1 cluster might only be observed when ensembles of a particular size are activated. We assessed this relationship by calculating the median number of units across ensembles in each cortical state cluster. Ensemble size differed across clusters (Kruskal-Wallis test, p = 0.0029, = 0.9608, = 13.97), but only between clusters 1 and 3 (**Figure 7C**). These results demonstrate that, while cluster 1 and 3 differ, there is no systematic relationship between the size of an ensemble and cortical state. Overall, our results demonstrate that cortical state depends on which specific ensembles are active, rather than simply an overall increase in the number of active single units or their firing strength.

### Activating a larger pool of LC ensembles results in a more homogeneous cortical state

These data clearly demonstrate a relationship between distinct LC ensembles and different cortical states. This finding stands in marked contrast to the stereotypical activated state evoked by stimulation of the LC, which evokes *en masse* spiking by LC neurons. Therefore, we predicted that when LC ensembles are co-active (i.e., more of the LC neurons are activated synchronously and LC population activity becomes more similar to stimulation-evoked *en masse* LC activation), the associated cortical state should become more homogenous to the activated state observed in studies that stimulated the LC. We took advantage of our observation that pairs of LC ensembles can sometimes become co-active (**Figure 5B**). We assessed the cortical LFP spectra, as in **Figure 6**, but triggered cortical spectrograms only on coactivation times of ensemble-pairs. A total of 199 ensemble-pairs had a significant zero-lag positive interactions in their cross-correlograms, which indicates co-activation. In contrast with the four heterogenous cortical states observed during activation of individual LC ensembles, k-means clustering now revealed only two types of cortical power spectra at the time points when ensemble-pairs were synchronously co-activated (**Figure 8**). One cluster is the stereotypical activated cortical state (cluster 2, 103 of 199 ensemble-pairs) and the other cluster is a homogenous decrease in spectral power (cluster 1, 96 of 199 ensemble-pairs). Therefore, when multiple LC ensembles are co-active, such that LC population activity becomes more similar to *en masse* LC activation, the modulation of cortical state is more homogenous and similar to LC stimulation-evoked cortical state changes (10–13).

**Figure 8.**
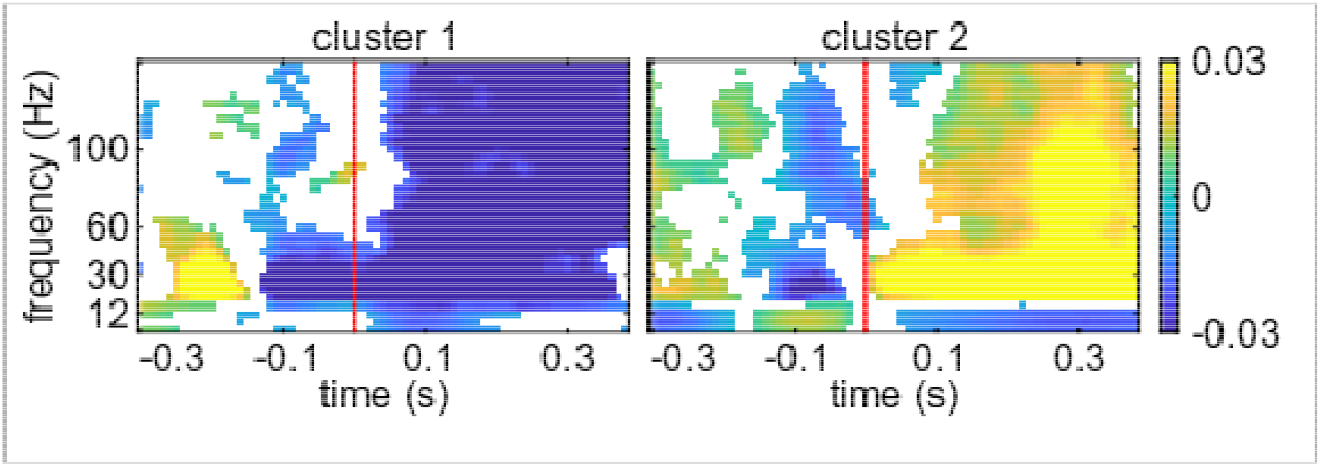
Synchronous coactivations of LC ensemble-pairs are associated with more homogeneous changes in cortical LFP power spectra. The LFP power spectra were triggered on the co-activation times of LC ensemble-pairs that had significant zero-lag cross-correlations. The resulting spectra clustered into 2 spectral types. Each plot shows the average spectrogram across all ensemble-pairs associated with each type of spectrum. Only significant changes in the power spectrum are plotted in color; non-significant modulations are white.

## Discussion

Cortical states can vary over a wide range and have been shown to be in a tight relationship with many functions that are relevant to psychiatric disorders, such as sleep, arousal, perceptual ability, and reaction times. It is thus no surprise that there have been long-standing efforts to understand the neural factors contributing to cortical state fluctuations (1–3). However, the self-organized neuronal interactions that control cortical states remain largely elusive. Blocking out the effect of the external world (e.g., slow wave sleep or anesthesia) has proven to be a successful approach for dissecting the spontaneous brain-internal neuronal interactions that control cortical state (30). These approaches have been used to demonstrate that the LC evokes transitions to, and maintenance of, a single and unitary activated state in the cortex (10–13). However, rather than studying the spontaneous emergence of cortical states due to LC activity, these studies have used electrical or optogenetic stimulation of the LC, which evokes *en masse* LC population activity.

Here, we considered the effect of spontaneously occurring events of LC ensemble activations. We demonstrate for the first time that spontaneous LC population activity consists of multiple, discrete neuronal ensembles each with their own nuanced time course of activity by applying NMF, an emerging computational methodology. Using synthetic ground-truth spike trains with nuanced patterns of ensemble activity, we illustrate that the time-averaged correlation analysis and graph theory detection of communities used in prior work (20) cannot resolve the identity of individual ensembles. While the first large-scale single unit recordings in the LC did successfully demonstrate with time-averaged correlation analysis that LC populations did not fire *en masse*, it was insufficient to demonstrate whether the LC contains ensembles.

Using NMF to define the neurons in distinct ensembles and the time courses over which the ensembles were active, we could study for the first time the self-organized spontaneous interactions between LC neurons and cortical states (25, 31) and therefore contribute to long-standing efforts to understand the how cortical state might be generated (1–3). The LCET-LFP analysis demonstrated that ongoing cortical state differs depending upon which LC ensemble is activated. Importantly, these findings establish that LC ensembles do not simply evoke a stereotypical activated state in the cortex. Instead, the nuanced, temporally diverse and largely non-overlapping nature of spontaneous LC ensemble activations correspond to a diversity of cortical states. Equally importantly, in the relatively rare cases when LC ensemble-pairs are coactivated synchronously, which is a situation more similar to the highly synchronized and *en masse* LC population activation driven by LC stimulation (10–13), the diverse set of cortical states collapses toward the stereotypical activated state that has been uniformly observed across these prior studies using LC stimulation.

### Potential neurophysiological causes of the diversity in cortical state

Neuromodulation of different forebrain regions may alter the self-organized brain-internal neuronal interactions that produce various cortical states. LC neurons are broadly-projecting, but also have localized projections to the forebrain and release a range of neurotransmitters (32, 33); therefore, LC ensembles that project to different forebrain neuronal networks could affect how those networks self-organize cortical states. When considering how distinct LC ensembles could promote different cortical states, two potential factors for future study are the diversity of ensemble neurochemical make-up and/or its projection profile. Given that the region in which we assessed cortical state (area 24a) receives projections from approximately 61 to 65% of LC neurons in the rat (34, 35), it seems likely that most of the ensembles project to area 24a and they should, therefore, produce a similar state change. Our finding to the contrary could be explained by the possibility that the neurochemical make-up of the LC neurons differs across ensembles and results in cortical state diversity. Another possibility is that, in spite of most ensembles presumably sharing area 24a as a projection target, it is the other targets that are potentially not shared across ensembles, which leads to LC ensemble-specific cortical states in 24a. According to this forebrain ‘network’ perspective, LC ensembles associated with different cortical states could have divergent axon collaterals which enable the ensembles to modulate distinct forebrain neuronal networks that are associated with different cortical states.

### Nuanced activation patterns of diverse LC ensembles enable greater diversity in neuromodulatory functions

Behavioral and mental states fluctuate widely from moment-to-moment and it has been known since the advent of EEG recordings that such diverse cognitive-behavioral states are associated with a multitude of cortical states (1–3). The brain-internal interactions that generate this large state-space are still largely unknown. LC neurons were classically thought to modulate cortical and thalamic neuronal excitability level using noradrenergic “tone” (4–7), whereas the neuronal interactions that produce the cortical state are contained within the cortex and thalamus (36, 37). According to this standard view, the role of the LC has been to modulate or predispose cortico-thalamic circuits toward the activated cortical state (and predispose the organism toward wakefulness), but the actual neuronal interactions that select cortical state are between cortex and thalamus. However, this standard view of cortical state generation was developed using methods that artificially activated the LC neuronal population *en masse* using external stimulation. Here, we used LCET-LFP to study the spontaneously self-organized neuronal interactions which are internal to the brain to reveal that, in contrast to this classical thinking, distinct LC ensembles can promote diverse cortical states. Each ensemble in this a small population of ∼1,600 brainstem noradrenergic neurons may individually be a key player in selecting ongoing cortical state from a multitude of possibilities. Thus, our findings shift the role of the LC from ‘modulator / promoter’ of a single cortical state toward a ‘selector / controller’ from a large sub-set of cortical states.

Our results imply that a single brainstem nucleus can perform different neuromodulatory functions by simply changing the groups of neurons that fire. Critically, these neuromodulatory dynamics can rapidly change on timescales relevant to flexible and ever-changing cognitive-behavioral states. The distinct LC ensembles with nuanced activation dynamics shown here substantially enrich the kind of functions that are currently attributed to brainstem nuclei.

## Materials and Methods

### Recording procedure and signal acquisition

All experiments were carried out with approval from the local authorities and in compliance with the German Law for the Protection of Animals in experimental research and the European Community Guidelines for the Care and Use of Laboratory Animals. Male Sprague-Dawley rats (350 - 450 g) were used (specific pathogen free, Charles River Laboratories, Sulzfeld, Germany). They were pair housed. Experiments were carried out during the active period of the rats, which were housed on a light cycle of 08:00 to 20:00 darkness. A sub-set of the data were collected from rats used in a prior study (20).

Neuronal recordings were made under urethane anesthesia, a widely-used model for studying cortical state transitions evoked by LC stimulation (10, 38). To date, recordings of many LC single units simultaneously in any awake organism with multi-electrode probes has been an intractable problem due to brainstem movement associated with body movement, thus necessitating the use of anesthesia to investigate the relationship between LC ensemble activity and cortical state. Rats were anesthetized using an intra-peritoneal (i.p.) injection of urethane at a dose of 1.5 g/kg body weight (Sigma-Aldrich, U2500). Surgical procedures were as described in prior work (20). Electrodes targeted the LC and the prelimbic division of the medial prefrontal cortex. The coordinates for LC were 4.0 mm posterior from lambda, 1.2 mm lateral from lambda, and approximately 6.0 mm ventral from the brain surface (implanted at a 15 deg posterior angle). The coordinates for the cortex were 3.0 mm anterior and 0.8 mm lateral from bregma and 3.0 mm ventral from the brain surface. The LC electrode was targeted based on standard electrophysiological criteria (20). At the end of the recording, we administered clonidine (0.05 mg/kg) i.p. (Sigma-Aldrich, product identification: C7897) to confirm cessation of noradrenergic neuron spiking. We also verified LC targeting in most experiments using histological examination of coronal sections (50 um thick) that were stained for Cresyl violet or a DAB and horse radish peroxidase reaction with hydrogen peroxide to visualize an antibody against tyrosine hydroxylase, as shown in prior work (20).

The LC was recorded using a multi-channel silicon probe (NeuroNexus, Model: A1×32-Poly3-10mm-25s-177-A32). The impedance of the electrodes was ∼1.0 to 2.0 MOhm. Cortical local field potentials were recorded using a single tungsten electrode with an impedance of 200 – 800 kOhm (FHC). A chlorided silver wire inserted into the neck muscle was used as a ground. Electrodes were connected to a pre-amplifier (in-house constructed) via low noise cables. Analog signals were amplified (by 2000 for LC and 500 for cortex) and filtered (8 kHz low pass, DC high pass) using an Alpha-Omega multi-channel processor (Alpha-Omega, Model: MPC Plus). Signals were then digitized at 24 kHz using a data acquisition device (CED, Model: Power1401mkII).

### NMF decomposition of population spike trains into coactive ensembles

We used non-negative matrix factorization (NMF) (39) to decompose a matrix of the spike counts of all simultaneously recorded single units across time intervals. NMF linearly decomposes the matrix of the spike counts of the population of single units at each time interval as a sum across a set of non-negative basis functions (modules) using non-negative coefficients (23, 24, 39). The non-negativity constraint is useful for obtaining sparse representations and it is particular suitable for decomposing population spike count at different time intervals, which are always non-negative. Previous work has shown that the NMF of population spike trains provides a robust decomposition whose basis functions can be biologically interpreted as a set of the firing patterns of the single units that are coactive (i.e., an ensemble) and the coefficients quantify the relative strength of recruitment of each ensemble firing pattern at any given time (23).

We employed an NMF decomposition that we have previously termed “space only NMF” because it decomposes the population firing patterns across single units at each time interval (23):

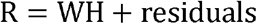

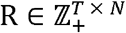 is the data matrix containing the spike counts of each of N single units binned into T time bins (with being the index of each time bin). 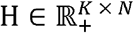 is the matrix containing the basis function, which has K spatial modules. Each module captures a different pattern of coactivity of the single units and can, therefore, be used to identify which neurons are active together and thus form ensembles. 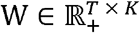 is the matrix containing the activation coefficients that describe the strength of recruitment of each module (and thus of each ensemble of coactive neurons) at each time interval. The residuals express the error in the reconstruction of the original population spike train matrix. We computed the decomposition using the multiplicative update rule to minimize the Frobenius norm between the original and the reconstructed data (39). Note that the use of the Frobenius norm assumes that the residuals have a Gaussian white noise structure.

One free parameter of the analysis is the temporal resolution of the time binning, ΔT. We binned spike counts at ΔT = 100 msec. The time resolution was selected based on our previous work reporting that pairs of LC single units are predominantly synchronized on a timescale of approximately 100 msec or less (20). We also used ranges of ΔT from several tens of msec to a few hundreds of msec and found that shorter bins (<=20 msec) and longer bins (> 1s), which our prior work suggests would be outside the range of LC single unit synchrony, tend to artificially identify either many modules each containing only one single active unit or one large ensemble containing all single units, respectively.

The second free parameter of the NMF analysis is the number of different modules,K, which were chosen for computing the decomposition. Following established procedures (23, 24), we chose K for each rat by computing the amount of the variance explained by the decomposition when varying K from its minimum possible value (one) to its maximum possible value (the number of simultaneously-recorded single units). An elbow in this plot indicates a point of diminishing returns for including more modules. We thus chose the number of modules as the smallest K in the elbow region of this curve for which the decomposition reconstructed at least 60% of the variance of the original spike train data. Given that the NMF decomposition may have local maxima in the variance explained (or equivalently local minima in error reconstruction), after selecting K, we repeated the decomposition five times using this K and used randomly chosen initialization conditions on each repetition. The selected K was used if all solutions had a high degree of stability across these five random initializations. Stability was assessed by checking the repeatability of clustering in comparison to randomly assigning single units to ensembles. The degree of stability was computed as follows. We hard clustered the data to assign each single unit to one and only one ensemble by dividing each column of H by its maximum and removing the values below 1. From these data we then measured the stability across the five decompositions using the Rand Index (40). We compared the average of the Rand index for each animal with 100 repetitions of the five random clustering. The average Rand Index was always greater than the top 5% of the distribution of mean Rand Indices resulting from random clustering. Therefore, NMF decomposition produced meaningful and repeatable ensembles. Among those random initializations, the final decomposition reported in the analyses was chosen as the one leading to the maximum variance explained.

The modules detected by NMF provide a pattern of coactivation of different single units and the activation coefficients measure the strength of recruitment of each module at any given time. From these data, we used a threshold-crossing of the coefficients to define when ensembles were active and which single units were active in the ensemble. In order to perform the thresholding, we first normalized the columns of H to the minimum and maximum and then set a threshold based on the distribution of coefficients. Single units within a module were defined as an ensemble of coactive single units if their corresponding element of H crossed the initial big peak of the histogram of the distribution of coefficient values for that rat (which usually corresponded to 95^th^ percentile). Coefficients below this value were set to zero and values above the threshold were set to one. In the resulting binary version of the matrix, H, a value of 1 represented spatial modules corresponding to a single unit belonging to an ensemble.

The columns of the W matrix correspond to a set of activation coefficients representing the strength of recruitment of each module at any given time interval. We thresholded these continuous values into binary values using the same method explained above for the spatial modules. The binary version of the matrix, W, hereafter referred to as “activation coefficient matrix,” was used to determine whether an ensemble is active or not in each time bin.

### The evaluation of physical clustering of ensembles according to location on the recording array

To assess whether single units within an ensemble tended to cluster on the recording array, we measured the pairwise distance between the units within each ensemble. The location of each unit was assigned to the electrodes on which the maximal waveform was recorded. Euclidian pairwise distances of the units inside each ensemble were calculated.

### Spike train simulation

In order to compare the NMF algorithm with previous work that used graph-theoretic community detection of sub-sets of coactive neurons from the time-averaged correlations between LC cell-pairs (20), we generated three different sets of spike trains. Each set of spike trains had unique ensemble dynamics as a ground-truth. Spike trains were generated using Poisson Process, with time varying rate every 100 msec. The baseline rate was randomly sampled from a Gaussian distribution with a mean of 1 and standard deviation of 0.1. For the periods of coactivation, the rate was increased by a signal to noise ratio sampled from a Gaussian with a mean of 1.5 and standard deviation of 0.1. After the generation of the spike trains, we binned and counted the spikes every 100 msec. This matrix was fed to NMF for extraction of spatial and temporal modules. Pairwise spike count correlation was calculated using Pearson correlation on the spike count matrix. A graph was made on the correlation matrix with each unit as a node and the links with the nodes was drawn only for significant correlations. The significance of the correlation was assessed by comparing to 1000 surrogate correlations generated by shuffling spike counts randomly in time. The graph was then used as an input for the Louvain community detection as described in prior work (20).

### The assignment of single unit types in the ensembles

Single unit type was defined by waveform duration, as in prior work (20). We determined if single units of the same type were more likely than chance to belong to an ensemble by computing the exact probability of having ensembles of the same single unit type under the null hypothesis of random assignment. These probabilities were computed by the means of repetition of random sampling (assembling) without replacement. The number of units in the sample was fixed to the number of single units in the ensemble. The number of repetitions for each rat was the number of ensembles that were empirically found by NMF to consist of only one type of single unit.

### Calculation of cross-correlograms between pairs of ensembles

Interactions between pairs of ensembles were measured using cross-correlograms between their time-dependent activation coefficients. Cross-correlograms were calculated in a window of 2000 msec with a bin size of 100 msec. The cross-correlograms were compared to 1000 surrogate cross-correlograms by jittering the activation times uniformly between ±1000 msec. Significant excitatory interactions were those that had cross-correlogram bins which crossed the upper 1% of pairwise coincidental activations observed in the surrogate data.

A synchrony index was used to measure the degree of synchrony between the ensemble pairs that show significant zero-lag synchronous activation. We calculated the synchrony index by dividing the number of co-activations of the two ensembles by the sum of total number of activation of each of the ensembles.

### Single unit spike rate PETHs and PETH clustering

The Peri-Event Time Histogram (PETH) of the spike times of single units inside and outside of an ensemble were aligned to events (at t = 0 msec), which were the ensemble activation times. We examined spike rate during a window from 100 msec before up to 400 msec after the ensemble activation times and used 1 msec bins. For each single unit, we calculated the average spike rate across activation events as though they were different “trials”. PETHs were smoothed with a Gaussian kernel (10 msec width). The PETH for each ensemble was obtained by averaging PETHs across all single units that were active in the ensemble.

PETH clustering was done in two steps. First, the dimensionality of the original PETHs in time was reduced using the Principle Component Analyses (41). Two dimensions explained more than 95% of the variance in the original data. After visualizing the data in the 2 dimensions we observed 3 non-circular masses of data. Therefore, we clustered the data in 3 groups using a Gaussian Mixture Model (GMM) (42). The GMM was calculated with 3 repetitions and full covariances.

### LC ensemble-triggered average LFP spectral modulation

We investigated the relation between the activation of LC ensembles and cortical state by triggering cortical LFP spectrograms on the timing of ensemble activation events. Spectra were computed using the multitaper method implemented in Chronux toolbox with 3 tapers and time bandwidth product of 5 (43). Short-time Fourier transforms were computed in a 10 msec moving window with a duration of 200 msec. The resulting spectral resolution was ∼4 Hz and the temporal resolution was 10 msec. We then averaged the resulting event-triggered spectra across all detected activation events separately for each ensemble.

From the averaged spectra, we computed an ensemble activation-triggered spectral modulation that characterized the effects of LC ensemble activation on the cortical LFP power spectrum. The spectral modulation was calculated as follows. We first averaged the spectrogram in time at each frequency for the baseline duration (400 msec before the ensemble being active) and then subtracted the baseline averaged spectrogram from the original spectrogram at each time step and divided by their sum. This quantity varies between -1 to 1 for each time *t* and frequency *f* and describes the average change in cortical LFP power around the time of ensemble activation.

### Spectrogram clustering

The set of so obtained ensemble activation-triggered spectral modulations were clustered, in order to assess the diversity of LC ensemble activation-triggered cortical states. The clustering was performed using the k-means algorithm (44). The k-means algorithm requires specifying a choice for the number of possible clusters and for the mathematical function used to compute the distance between the different spectrograms. We tried various definitions of distance functions (Pearson Correlation, Euclidean distance, cosine, and cityblock), and we chose Pearson correlation as distance function because it gave higher averaged silhouette values (45) (i.e., cleaner clustering). We clustered the spectral modulation into k=4 clusters. This number of clusters was selected because it corresponded to the elbow point (defined as the first point in which the error drops below 5%) of the curve quantifying the normalized clustering error (error divided by the maximum error) as a function of the selected number of clusters. The error in the k-means clustering is computed as sum of the distances of each data point to their respective cluster centroid. We assessed the significance of the clustered spectral modulations at each time and frequency by pooling the spectral modulations of all ensembles in each cluster by comparing the median of the population at each point against zero using Wilcoxon signed rank test (5% significance level). The p-values were corrected for multiple comparisons using Benjamini’s & Hochberg’s method for false discovery rate (46).

The above analysis was done taking for clustering all spectral modulations obtained in correspondence of a detected activation of one or more ensembles. We performed a further control analyses in which we clustered only the subset of the spectral modulations during coactivation of ensemble-pairs. The clustering procedure for this control analysis was identical to the one reported above but selected a number of clusters (corresponding to the elbow point of the error curve) equal to 2 clusters.

## Acknowledgements

This research was supported by the Helsinki Institute of Life Science (NKT) and the Department of Physiology of Cognitive Processes in the Max Planck Institute for Biological Cybernetics (IZ, NKL, NKT). SP was supported by the Fondazione Caritro and by a SFARI explorer grant (Grant no. 602849).

## Author Contributions

Conceptualization: NKT, SP; Methodology: NKT, SN, SP; Formal analysis: SN; Investigation: IZ, NKT; Resources: NKL, SP; Writing: NKT, SN, SP; Visualization: SN; Supervision: NKT, SP; Funding acquisition: NKL, NKT, SP.

## Declaration of Competing Interests

The authors declare no competing interests.

## Supplemental information

**Figure S1.**
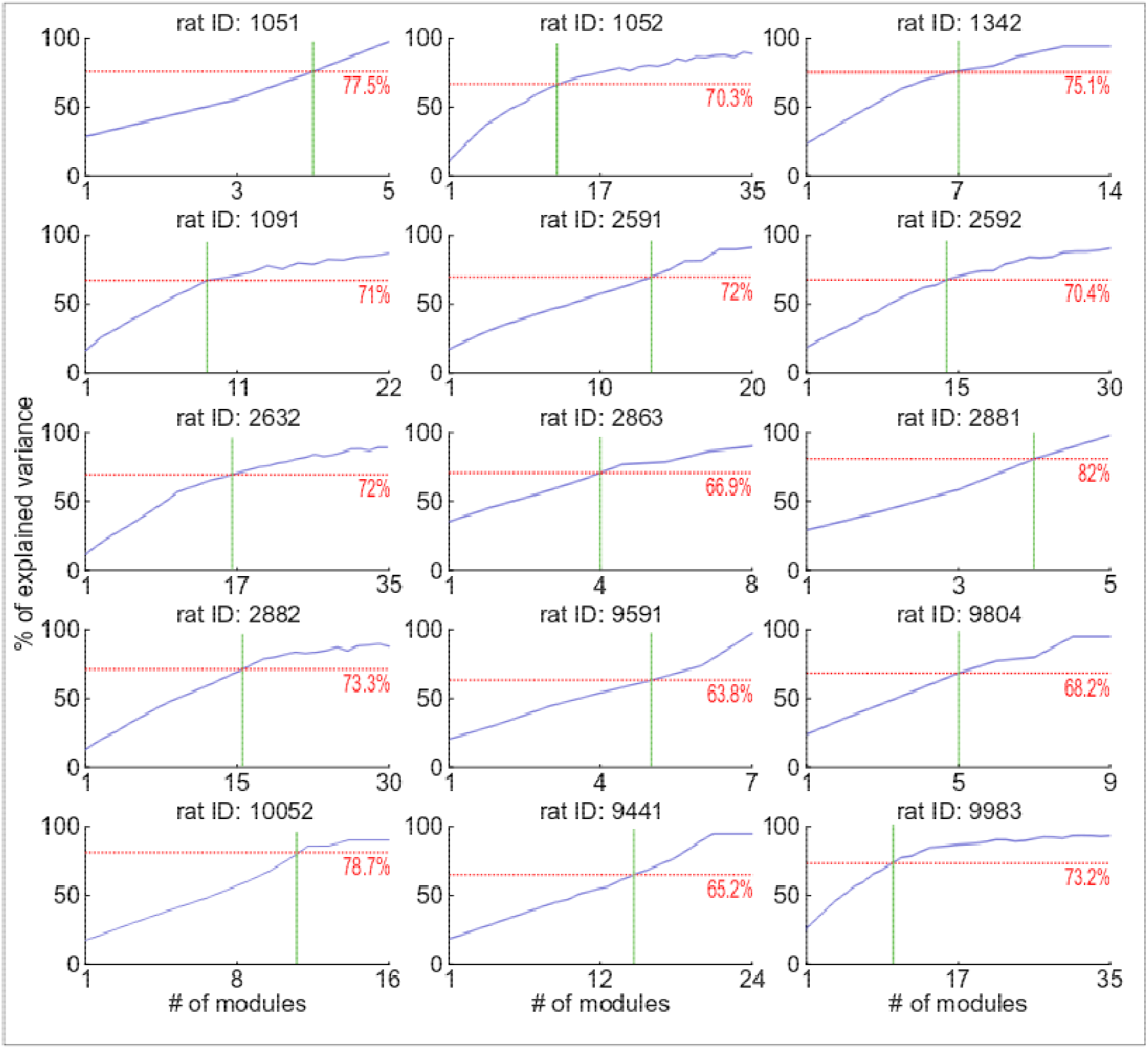
Data underlying the choice of the optimal number of modules (K) in each rat. Each panel depicts the percentage of explained variance versus the number of the modules for each rat. Solid green lines show the number of selected modules based on the criteria of first elbow after at least 60% of variance is explained. The dotted red lines show the amount of the explained variance at the selected number of modules.

**Figure S2.**
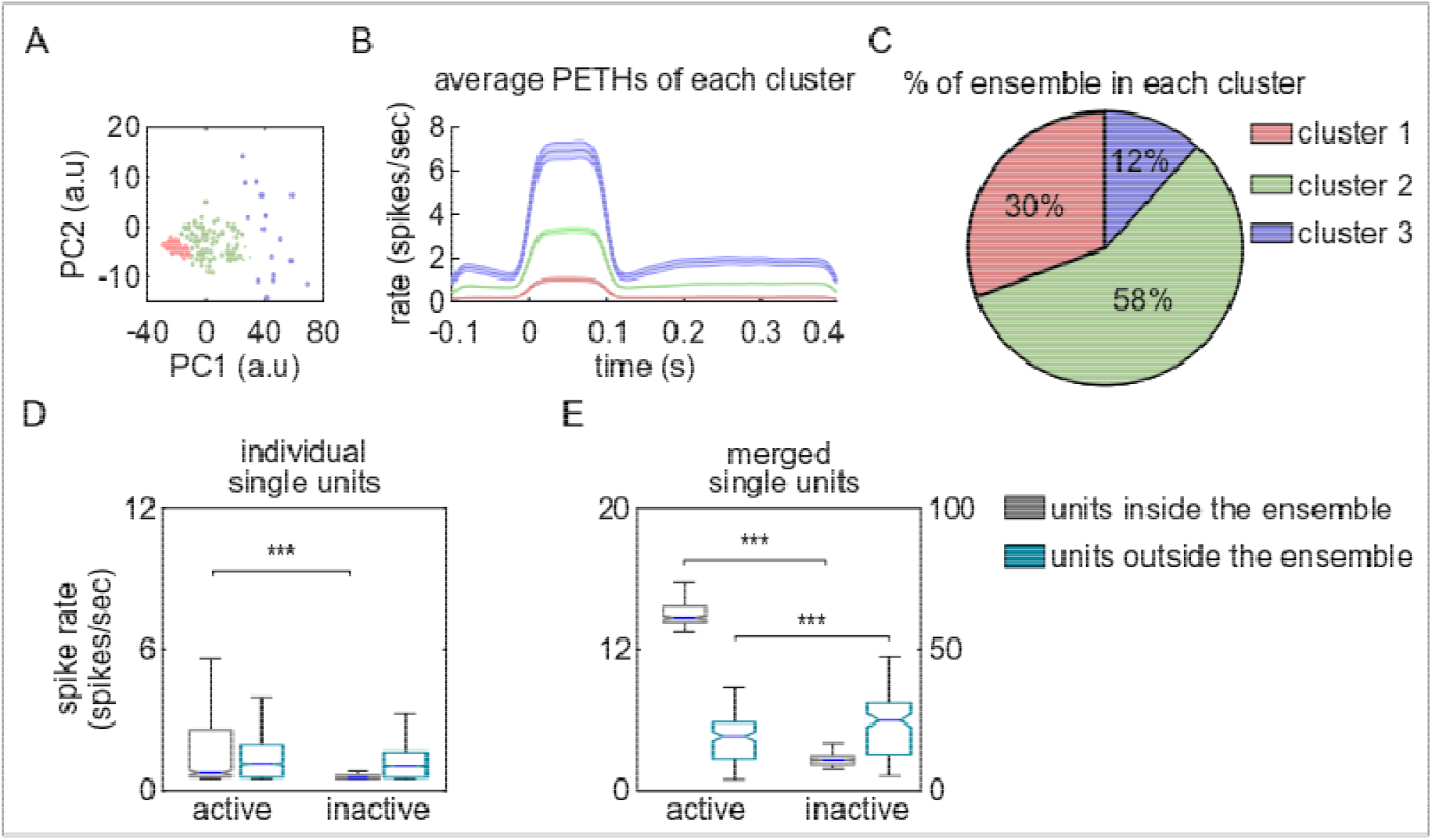
LC ensembles are characterized by different magnitudes of change in spike rate. **(A)** The scatter plot shows the projections of the PETHs into two dimensions (PC1, PC2) using PCA. The first two principal components explained more than 95% of the variance. Three non-circular masses were clustered using GMM. Data points falling into each cluster are color coded separately. **(B)** Average PETHs of the ensembles in the same cluster. The zero time on the x-axis is the ensemble active time. The PETHs of all ensembles were grouped into 3 clusters that increased their firing rate to different degrees. **(C)** The pie chart illustrates the percentage of ensembles in each PETH cluster. Most ensembles had a medium (green) or low (red) magnitude increase in single unit spike rate. **(D, E)** The box plots show the distribution of the spike rates for the single units inside the ensemble (gray) and outside the ensemble (light green). The result is shown separately for individual single units (C) and the spike trains merged across single units (D). The spike rate was calculated as the average of all ensemble activation events combined across all single units in the ensemble (i.e., in Figure 1B and 6A spike rasters, all events of different colors were averaged). Spike rate increased when the ensemble was in an active state for both single units and multi-unit activity. Additionally, when the ensemble was inactive, multi-unit activity outside of the ensemble was higher presumably due to units spiking in other ensembles.

**Figure S3.**
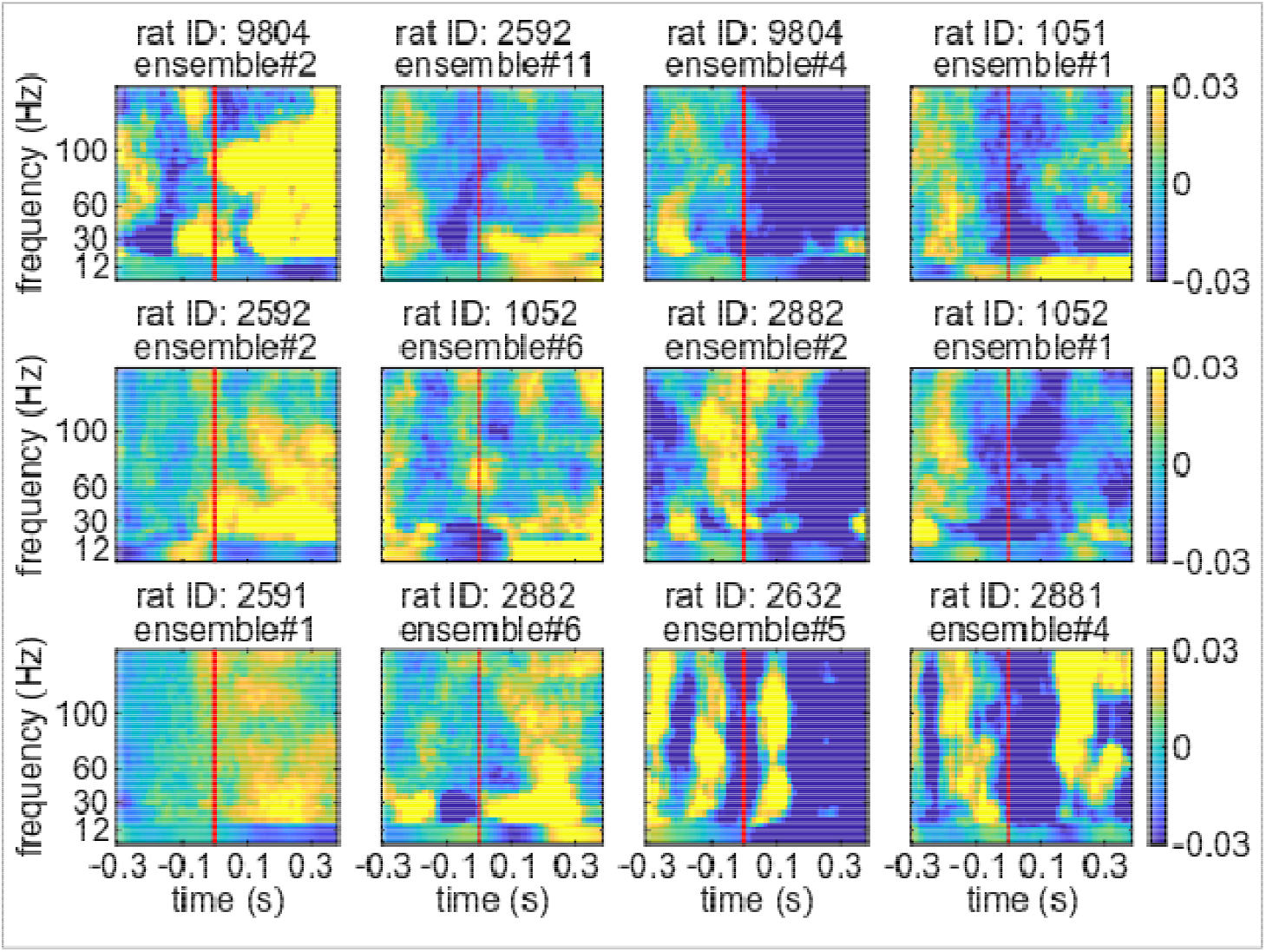
Examples of LC ensemble activation-triggered cortical LFP power spectra. Examples from 12 different LC ensembles illustrate the diverse cortical states which occur around the time of ensemble activation. The examples are shown in 4 columns, each of which indicates a specific trend in the spectra corresponding to the clusters shown in the Figure 5 in the Main Text.

**Figure S4.**
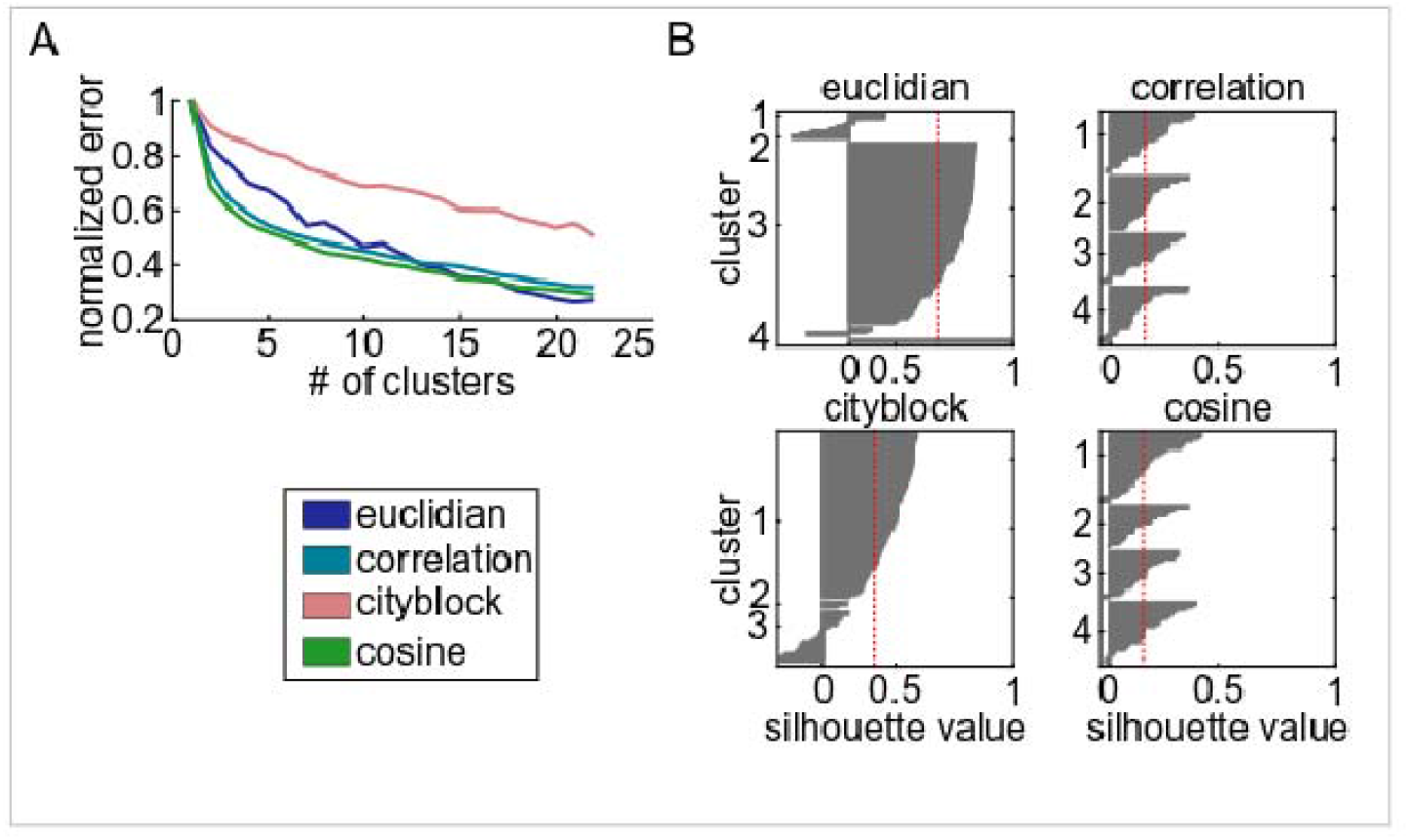
The result of analyses supporting the determination of the best criteria for spectral clustering. (A) The normalized error (error divided by the maximum error) of the k-means clustering of the ensemble activation-triggered spectra versus the number of clusters. Four different distance measures were assessed and each is plotted in a different color. (B) Each panel shows the result of the silhouette analyses on the chosen number of clusters for four different distance measures. The optimal distance was selected based on both the uniformity in each cluster (the width of the bar plots) and the average silhouette value (the dashed red line).

## Notes

### Competing Interest Statement

The authors have declared no competing interest.

### Summary of Updates

A new figure (Fig. 3) was added to demonstrate that NMF detects the composition of LC ensembles and their activation times, whereas graph-theoretic time-averaged pairwise correlations used in prior work does not. Additionally, the writing has been modified throughout to emphasise how the present study relates to prior work.

